# Complete loss of RelA and SpoT homologs in *Arabidopsis* reveals the importance of the plastidial stringent response in the interplay between chloroplast metabolism and plant defense response

**DOI:** 10.1101/2022.09.20.508797

**Authors:** Masataka Inazu, Takanari Nemoto, Sae Suzuki, Sumire Ono, Yuri Kanno, Mitsunori Seo, Akira Oikawa, Shinji Masuda

**Affiliations:** Department of Life Science and Technology, Tokyo Institute of Technology, Yokohama, 226-8501, Japan; RIKEN Center for Sustainable Resource Science, Yokohama 230-0045, Japan; Graduate School of Agriculture, Kyoto University, Uji 611-0011, Japan

**Keywords:** Chloroplast, Defense response, Photosynthesis, ppGpp, Salicylic acid, Stringent response

## Abstract

The highly phosphorylated nucleotide, guanosine tetraphosphate (ppGpp), functions as a secondary messenger in bacteria and chloroplasts. The accumulation of ppGpp alters plastidial gene expression and metabolism, which are required for proper photosynthetic regulation and robust plant growth. However, because four plastid-localized ppGpp synthases/hydrolases function redundantly, the impact of the loss of ppGpp-dependent stringent response on plant physiology remains unclear. We used the CRISPR/Cas9 technology to generate an *Arabidopsis thaliana* mutant lacking all four ppGpp synthases/hydrolases, and characterized its phenotype. The mutant showed 20-fold less ppGpp levels than the wild type (WT) under normal growth conditions, and exhibited leaf chlorosis and increased expression of defense-related genes as well as salicylic acid and jasmonate levels upon transition to nitrogen-starvation conditions. These results demonstrate that proper levels of ppGpp in plastids are required for controlling not only plastid metabolism but also phytohormone signaling, which is essential for plant defense.

## Introduction

Adaptation to fluctuating environment is critical for organisms. In bacteria, one of the environmental adaptation mechanisms is the stringent response, which was first discovered in *Escherichia coli* as an adaptive response to amino-acid starvation conditions and was characterized by a rapid reduction in rRNA and tRNA synthesis (Cashel, 1969). The stringent response is regulated by the hyperphosphorylated nucleotides guanosine 5′-triphosphate 3′-diphosphate and/or guanosine 5′-diphosphate 3′-diphosphate (ppGpp). In *E. coli*, two different enzymes, RelA and SpoT, respond to multiple stresses such as fatty acid deprivation and heat stress, in addition to nutrient starvation, by regulating endogenous ppGpp levels (Cashel *et al*., 1996; Potrykus & Cashel, 2008). The synthesized ppGpp controls transcription, translation, DNA replication, and various enzymatic activities to adapt to environmental stresses.

Owing to the vast amounts of publicly available genome sequence data, RelA-SpoT homologs (RSHs) have been identified in a wide range of eukaryotes including algae, metazoa, and higher plants, indicating that the ppGpp-dependent regulation of cellular activities is highly conserved among eukaryotes (van der Biezen *et al*., 2000; Givens *et al*., 2004; Tozawa *et al*., 2007; Masuda *et al*., 2008; Tozawa & Nomura, 2011; Ito *et al*., 2017, 2020, 2022; Imamura *et al*., 2018; Field, 2018; Avilan *et al*., 2019, 2021; Li *et al*., 2022). Plant RSH proteins have been extensively studied in recent years. In the model plant *Arabidopsis thaliana*, four RSH proteins, RSH1, RSH2, *RSH3*, and Ca^2+^-activated RSH (CRSH), have been identified to date (Tozawa & Nomura, 2011; Masuda, 2012; Field, 2018). All of these four RSHs are encoded by the nuclear genome, possess a chloroplast transit signal at their N-terminus, and localize to chloroplasts. The activities of RSH proteins are regulated at the transcriptional and posttranslational levels to coordinately control plastidial ppGpp levels (Mizusawa *et al*., 2008; Yamburenko *et al*., 2015; Sugliani *et al*., 2016; Ono *et al*., 2021), which transiently increase upon the light– dark transition, under various abiotic stresses, and upon treatment with phytohormones such as abscisic acid (ABA), jasmonic acid (JA), and ethylene (Takahashi *et al*., 2004; Ihara *et al*., 2015). The ppGpp nucleotide represses the transcription of some chloroplast genes and controls specific enzyme activities both *in vivo* and *in vitro* (Yamburenko *et al*., 2015; Maekawa *et al*., 2015; Sugliani *et al*., 2016; Ono *et al*., 2021; Avilan *et al*., 2021; Romand *et al*., 2022). Furthermore, the excessive accumulation of ppGpp due to *RSH3* overexpression potentially contributes to the adaption of plants to nitrogen deficiency (Maekawa *et al*., 2015; Honoki *et al*., 2018; Goto *et al*., 2022). These results indicate that the plastidial ppGpp plays a crucial role in various stress responses in plants.

However, given that *Arabidopsis* RSHs function redundantly to maintain an optimal level of ppGpp, the complete loss of RSHs has long been awaited to further reveal the functional roles of RSHs in the ppGpp-dependent plastidial stringent response. In this study, we successfully constructed *Arabidopsis* quadruple *rsh* null mutant that lacks all four RSHs, and investigated the impact of the complete loss of RSHs on the plant response to diverse environmental changes. Our results provide direct evidence supporting the role of plastidial ppGpp homeostasis in plant metabolism, photosynthesis, hormone signaling, and consequently the plant defense response.

## Results

### Isolation of the quadruple *rsh* mutant

Previous quadruple *rsh* mutants (QMai or QMaii), in which *CRSH* was knocked-down, were constructed using the RNA interference (RNAi) technique on the *rsh1 rsh2 rsh3* triple mutant (Sugliani *et al*., 2016). The QMai and QMaii mutants showed altered expression of defense-related genes (Abdelkefi *et al*., 2018; Romand *et al*., 2022), suggesting the importance of ppGpp-dependent plastidial stringent response for the host plant physiology. Nonetheless, both QMai and QMaii could accumulate ppGpp to a certain level (∼50% of that in the WT), perhaps because of residual CRSH activity (Sugliani *et al*., 2016). Therefore, we assumed that the isolation of another quadruple mutant, which completely lacks all four *RSH* genes, would be necessary to clearly evaluate the function and physiological importance of ppGpp. To isolate a new quadruple *rsh* null mutant, we used the CRISPR/Cas9-based genome editing technique (Fauser *et al*., 2014). A protospacer sequence was designed, as reported previously (Ono *et al*., 2021) to induce a double strand break in the first exon of *CRSH*, and the CRISPR/Cas9 construct was introduced into the *rsh1 rsh2 rsh3* triple mutant (Ono *et al*., 2021). Twenty independent T1 transgenic plants were selected, and one of the mutant lines (T3 generation) showed a single nucleotide insertion at the 99th nucleotide position of *CRSH* (Fig. 1a). This mutation was predicted to cause a frameshift in the first exon of *CRSH*, leading to a premature stop codon at the 48th amino-acid position of the CRSH protein. Western blot analysis using anti-CRSH antibody confirmed the absence of CRSH in four individuals (#1–4) of the mutant line at the T3 generation (Figs. 1b, c).

**Fig. 1.**
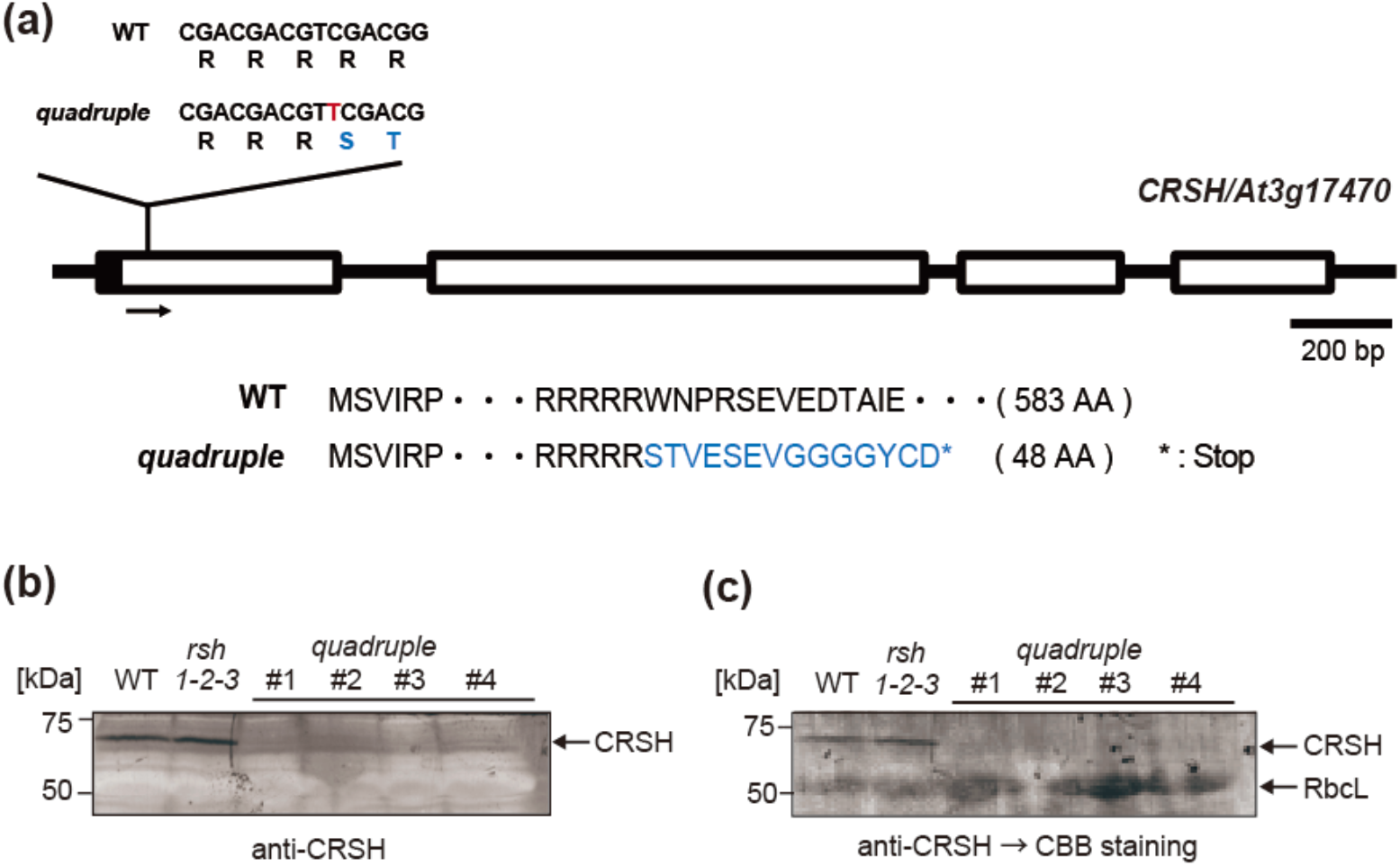
Construction of the *rsh* quadruple mutant. **(a)** Schematic of the exon–intron structure of *CRSH* (At3g17470). A single nucleotide insertion and mutated codons in the quadruple mutant are highlighted in red and blue, respectively. **(b)** Western blot analysis of proteins isolated from the WT, *rsh1 rsh2 rsh3* triple mutant (*rsh1-2-3*), and four quadruple mutant plants (#1–4). Proteins were immunodetected using an anti-CRSH antibody. **(c)** Membrane shown in **(b)** stained with Coomassie brilliant blue (CBB).

### Accumulation of ppGpp is decreased in the *quadruple* mutant

To investigate whether all RSHs are dysfunctional in the quadruple mutant, we quantified ppGpp levels in the quadruple mutant plants grown under continuous light conditions. A significant reduction in ppGpp level was observed in the quadruple mutant (8.7 ± 1.5 pmol g^−1^), which was ∼5% of the ppGpp amount in the WT (172.9 ± 15.6 pmol g^−1^) (Fig. 2a). Furthermore, WT plants showed a transient increment in the amount of ppGpp upon the light-to-dark transition, followed by a decline in ppGpp levels, reaching basal levels at 60 min after the light-to-dark transition (Fig. 2b), as reported previously (Ono *et al*., 2021). On the other hand, the quadruple mutant showed no ppGpp increment upon the light-to-dark transition (Fig. 2b). These results indicated that all four RSHs were dysfunctional in the quadruple mutant, but it was unclear how the ppGpp detected in the quadruple mutant was synthesized. Given that the *rsh1 rsh2 rsh3* triple mutant showed significantly higher ppGpp accumulation (∼10-fold) compared with the WT upon the light-to-dark transition (Ono *et al*., 2021), the data obtained here further supported the previous model, according to which the dark-induced accumulation of ppGpp is caused by the activation of CRSH due to the dark-induced increment in Ca^2+^ concentration in the chloroplast stroma (Ono *et al*., 2021).

**Fig. 2.**
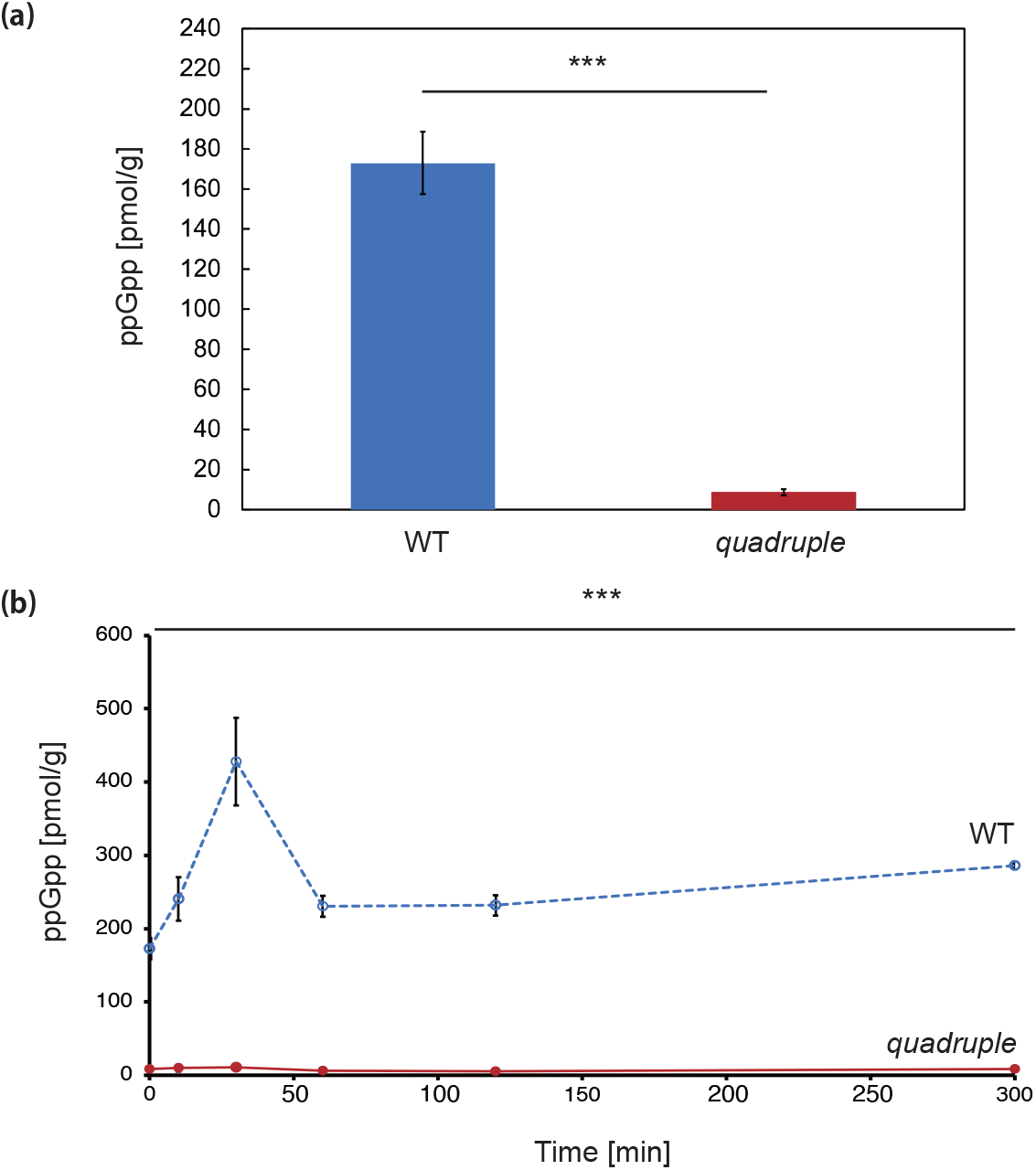
Accumulation of ppGpp in WT and quadruple mutant. **(a)** ppGpp levels of plants grown on +N medium under continuous light conditions for 14 days. **(b)** Accumulation of ppGpp in the WT and quadruple mutant during the light-to-dark transition. Plants were grown on +N medium under continuous light conditions. After 14 days, the plants were transferred to the dark at time 0, and shoots were harvested at the indicated time points for quantifying ppGpp. Data represent mean ± SD (*n* = 3). Asterisks indicate statistically significant differences between the quadruple mutant and WT (\*\*\**p* < 0.001; Student’s *t*-test).

### Visible phenotype of the quadruple *rsh* mutant

A recent study showed that overexpression of RSH1, which encodes the ppGpp hydrolase, in the *Arabidopsis* mutant decreased the ppGpp level by ∼50% and reduced the tolerance to nitrogen deprivation compared with the WT (Romand *et al*., 2022). This led us to hypothesize that ppGpp plays an important role in the plant adaptation to nitrogen starvation (−N). To test this hypothesis, we analyzed the response of the quadruple mutant to −N conditions. The WT and quadruple mutant plants were grown on nitrogen-rich (+N) medium for 14 days, and then transferred to +N and/or −N medium and grown for another 10 days. Comparison of ppGpp levels after the transition to −N or +N conditions showed that ppGpp levels in WT plants grown under −N conditions were 95.2 ± 7.8% of those under +N conditions (n = 3). Thus, the ppGpp levels showed no significant difference between −N and +N conditions. This result is consistent with a previous study, in which WT plants transiently accumulated ppGpp upon the transition to −N conditions; however, the amount of ppGpp returned to basal levels within 12 days (Romand *et al*., 2022). Although the quadruple mutant showed no visible phenotype under +N conditions compared with the WT, the mutant showed strong leaf chlorosis under −N conditions, as evident from the pale-green color of leaves, which was more severe in the mutant than in the WT (Fig. 3a). The chlorosis phenotype was accompanied by a reduction in fresh weight; the fresh weight of WT plants remained stable even after the transition to −N conditions, whereas that of quadruple mutant plants gradually decreased after the transition to −N conditions (Fig. 3b). Although the mean fresh weights of WT and quadruple mutant plants showed no significant difference before the transition to −N conditions, the fresh weight of the quadruple mutant was lower than that of WT at 12 days after the transition to −N conditions, indicating that a certain amount of ppGpp is required for the proper response to nitrogen deficiency.

**Fig. 3.**
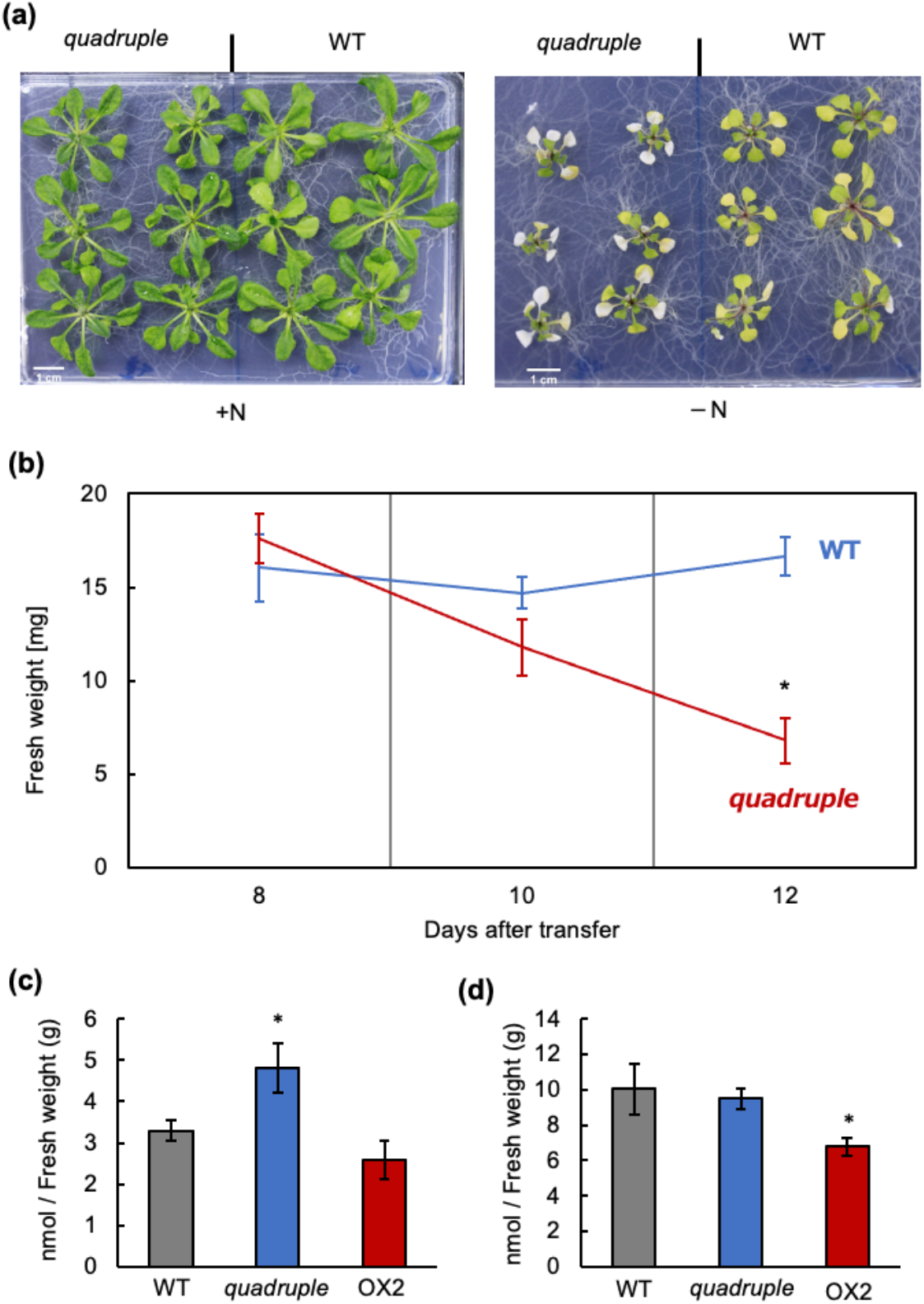
Phenotypes of quadruple mutant plants grown under −N conditions. **(a)** WT and quadruple mutant plants grown for 14 days under +N conditions, and then transferred to and grown for 10 days on +N or −N medium. **(b)** Changes in fresh weight after the transfer to −N medium. The shoot weight of each plant was measured separately. Data represent mean ± standard error (SE; *n* = 6–24). Asterisk indicates statistically significant differences between the quadruple mutant and WT (\**p* < 0.05; Student’s *t*-test). **(c, d)** Quantification of H_2_O_2_ in WT, quadruple mutant, and RSH3ox2 (OX2) mutant plants under +N **(c)** and −N **(d)** conditions. Plants were grown as in **(a)**. Data represent mean ± SD (*n* = 3). Asterisk indicates significant differences (\**p* < 0.05; Dunnett test).

Next, we quantified the Chl and Car contents of the quadruple mutant (Table 1). Under +N conditions, the total Chl *a* and Chl *b* contents showed difference between the quadruple mutant and the WT, although the Chl *a*/Chl *b* ratio was significantly higher in the quadruple mutant than in the WT. Under −N conditions, the total Chl and Car levels were significantly decreased in the quadruple mutant compared with the WT, and the Chl *a/*Chl *b* ratio was significantly higher in the quadruple mutant than in the WT, as observed under +N conditions. These results suggest that the alteration of basal ppGpp level affects pigment contents, especially under −N conditions.

**Table 1.** Pigment contents of wild-type (WT) and quadruple mutant *Arabidopsis* plants

| Genotype | Chl *a* | Chl *b* | Chl *a/b* | Total Chl | Total Car |
| --- | --- | --- | --- | --- | --- |
| +N |  |  |  |  |  |
| WT | 2.54<br>(±0.10) | 0.84<br>(±0.04) | 3.02<br>(±0.02) | 3.38<br>(±0.14) | 0.69<br>(±0.03) |
| *quadruple* | 2.31<br>(±0.09) | 0.74<br>(±0.03) | 3.12 *<br>(±0.02) | 3.06<br>(±0.13) | 0.66<br>(±0.05) |
| -N |  |  |  |  |  |
| WT | 0.52<br>(±0.01) | 0.18<br>(±0.01) | 2.92<br>(±0.04) | 0.70<br>(±0.02) | 0.29<br>(±0.01) |
| *quadruple* | 0.41 *<br>(±0.02) | 0.12 *<br>(±0.01) | 3.49 *<br>(±0.07) | 0.52 *<br>(±0.02) | 0.22 *<br>(±0.01) |

Plants were grown under continuous light on ½ MS medium for 14 days, and then grown on nitrogen-rich 1/2MS (+N) and/or nitrogen-starvation medium (–N) for 10 days. Chlorophyll (Chl) and carotenoid (Car) pigments were extracted from the shoots and quantified. The Chl and Car contents were normalized relative to the shoot fresh weight (FW), and expressed as mg mgFW<sup>–1</sup>. Data are represented as mean ± standard deviation (SD; *n* = 3). Asterisk indicates statistically significant differences between the quadruple mutant and WT (\**p* < 0.05; Student's *t*-test).

To further investigate the relationship between ppGpp levels and chlorosis phenotype, we quantified the level of H_2_O_2_, one of the reactive oxygen species (ROS), in the quadruple mutant, WT, and ppGpp-accumulating mutant RSH3ox2 (control; Maekawa *et al*., 2015). RSH3ox2 is an *RSH3* overexpression line in the *rsh2 rsh3* double mutant background, which shows ∼3-fold higher ppGpp levels than the WT under normal growth conditions (Maekawa *et al*., 2015). It should be noted that RSH3ox2 maintained the green color of leaves under −N conditions, whereas WT plants showed the senescence phenotype (Maekawa *et al*., 2015; Honoki *et al*., 2018; Goto *et al*., 2022). The quadruple mutant showed significantly higher H_2_O_2_ levels than the WT, although H_2_O_2_ levels were similar in the RSH3ox2 mutant and WT (Fig. 3c). Upon transition to −N conditions, all three genotypes showed an ∼3-fold increase in H_2_O_2_ level at 10 days after the transition to −N conditions (Fig. 3d). Under −N conditions, RSH3ox2 showed significantly lower H_2_O_2_ levels than the WT; on the other hand, no significant difference was observed in H_2_O_2_ levels between the WT and quadruple mutant (Fig. 3d). These results indicate that lower levels of ppGpp lead to the accumulation of ROS under +N conditions, and higher basal levels of ppGpp result in the reduction of ROS under −N conditions.

### Photosynthetic activity of the quadruple *rsh* mutant

To investigate the impact of significant reduction in ppGpp levels on photosynthetic regulation, we analyzed NPQ and Y(II) in WT and quadruple mutant plants grown under +N and −N conditions. Under +N conditions, the values of NPQ induction and Y(II) in the quadruple mutant were similar to those in the WT (Figs. 4a, b). Upon the transition to −N conditions, the value of NPQ increased drastically in the WT at the induction stage (1–2 min) compared with the +N conditions (Fig. 4a). On the other hand, compared with the WT, NPQ induction in the quadruple mutant was lower under −N conditions, and similar under +N conditions (Fig. 4a). Y(II) values both in the WT and *quadruple* mutant were not influenced by the transition to −N conditions (Fig. 4b), indicating that proper plastidial ppGpp level is required for NPQ induction under −N conditions. However, photosynthetic electron transfer was not influenced by −N conditions or by quadruple *rsh* mutations.

**Fig. 4.**
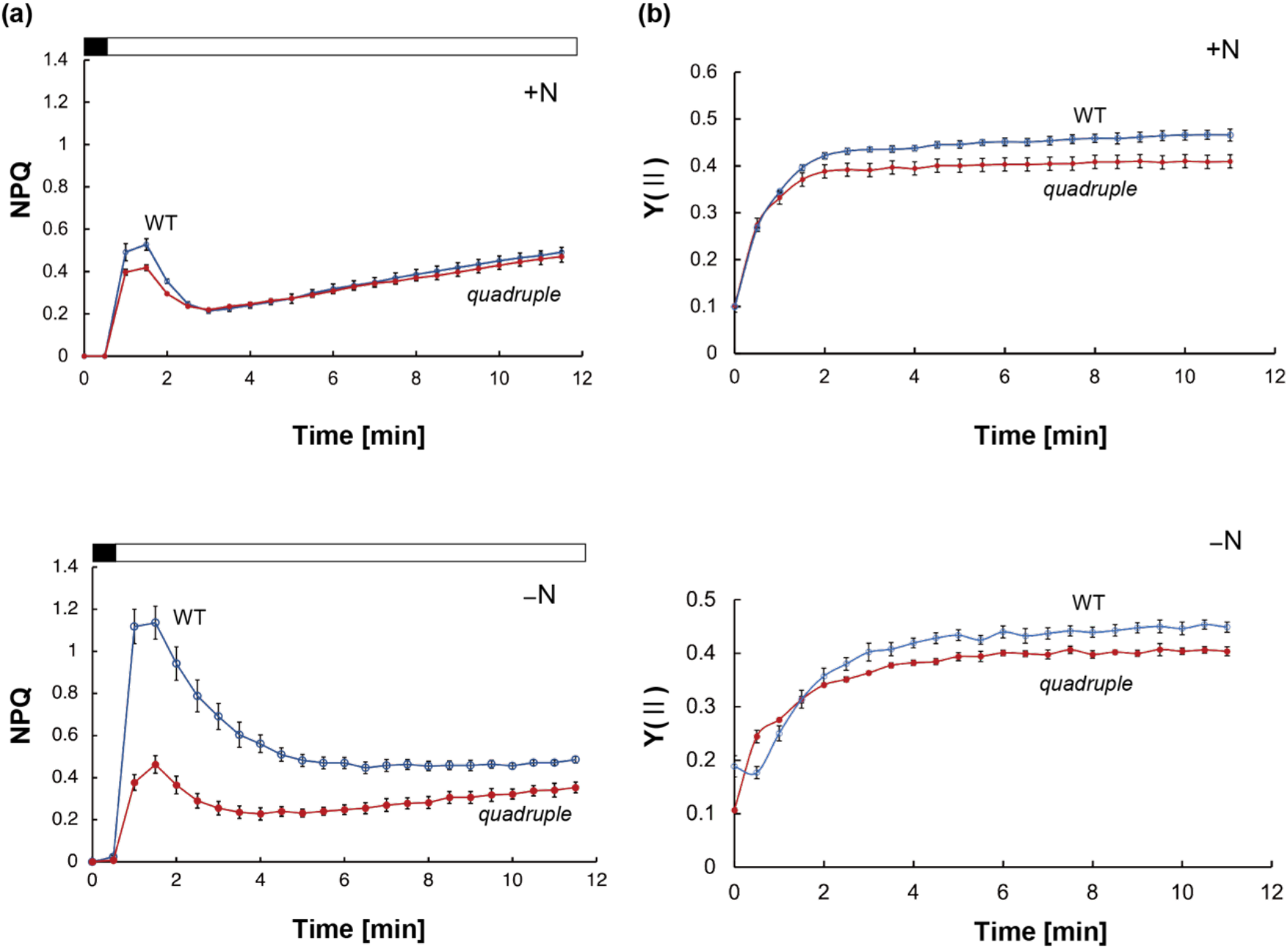
Measurement of photosynthetic activity. **(a)** NPQ; **(b)** Y (II). Plants grown under +N conditions for 14 days were transferred to +N or **−**N medium and grown for another 10 days. Data represent mean ± SE (*n* = 3). Measurements were taken under 54 µmol photons m^−2^ s^−1^ actinic light.

### Metabolite changes caused by reduced ppGpp levels

To investigate the influence of lowered basal levels of ppGpp on metabolic control, we performed metabolome analysis of WT and quadruple mutant plants, which were grown under normal +N conditions for 14 days and then under +N and/or −N conditions for 10 days. We were able to quantify 102 and 92 metabolites in plants grown under +N and −N conditions, respectively. To examine the effects of significant reduction in ppGpp on plant metabolism, we first performed global component analysis of the metabolome data by PLS-DA (see Materials and Methods for details). As shown in Fig. 5, all individual datasets were divided into four clusters within both component 1 and component 2, which reflect different genetic and growth conditions, respectively. Specifically, different growth conditions (+N and −N) were separated in component 1, whereas the genetic backgrounds of the WT and quadruple mutant were separated in component 2. Consequently, the separation between WT and quadruple mutant was relatively larger under −N conditions than under +N conditions. These results suggest that the function of RSH is required for regulating plant metabolism not only under −N conditions but also under +N conditions, and it is particularly required for metabolic changes induced by the transition from +N to −N conditions.

**Fig. 5.**
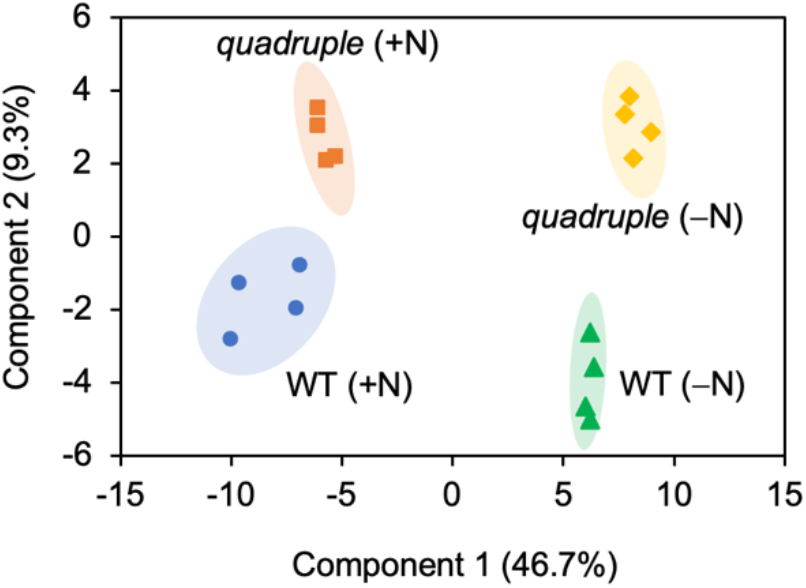
Global component analysis of metabolites. Plants grown for 14 days under +N conditions were transferred to +N or −N medium and grown for 10 days. Then, shoots (∼ 40 mg) of each plant were harvested separately and subjected to metabolome analysis. Component 1 and component 2 were obtained by the PLS-DA based on the metabolome data of WT and quadruple mutant plants grown under +N and −N conditions. Experiments were repeated four times, and the data from each experiment were plotted separately.

Next, we measured the level of each metabolite under +N and −N conditions. Under +N conditions, the quadruple mutant showed differences in some metabolite levels compared with the WT (Fig. 6a). Specifically, the amount of γ-glutamylcysteine, which is involved in nicotinic acid and glutathione metabolism, was higher in the quadruple mutant than in the WT. On the other hand, GSH and CysSG, which are involved in the metabolism of glutathione, shikimic acid in the shikimic acid pathway, succinate in the TCA cycle, and Gln in the glutamine synthetase/glutamine oxoglutarate aminotransferase (GS/GOGAT) cycle, were significantly lower in the quadruple mutant than in the WT.

**Fig. 6.**
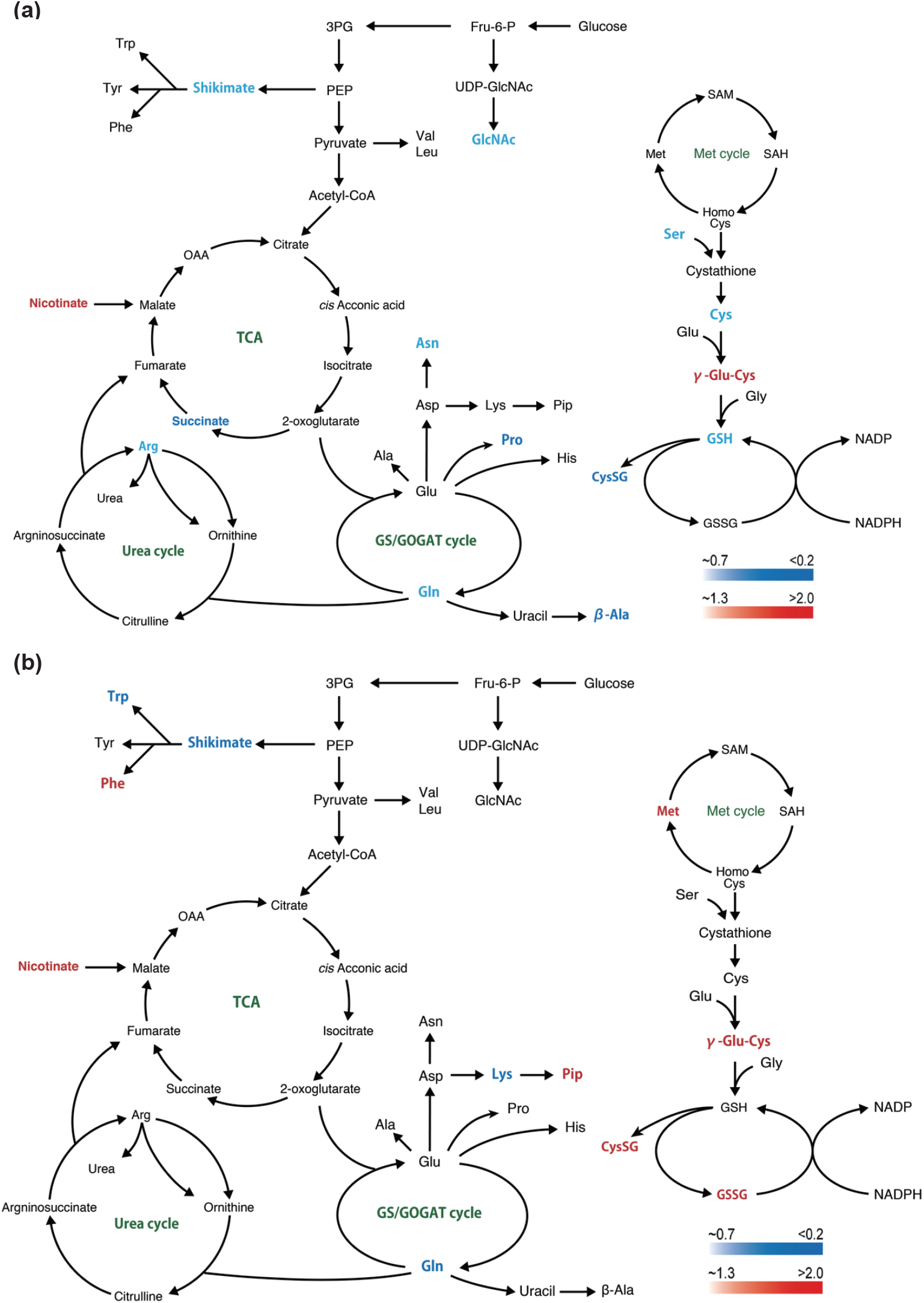
Statistically significant changes in metabolites between the quadruple mutant and WT plants under +N **(a)** and −N **(b)** conditions. Metabolome data used are the same as in Fig. 5. Significant changes in metabolite accumulation (*p* < 0.05, Student’s *t*-test, *n* = 4) are expressed along color gradients of red (increased) and blue (decreased). OAA, oxaloacetate; Pip, pipecolate; SAM, S-adenosylmethionine; SAH, S-adenosylhomocysteine.

Under −N conditions, the quadruple mutant showed a decline in the contents of shikimic acid and Gln, which are involved in the GS/GOGAT cycle, compared with the WT (Fig. 6b). On the other hand, nicotinic acid, CysSG, GSSG, and γ-Glu-Cys were increased approximately by 2-fold in the quadruple mutant compared with the WT. These results suggest that *rsh* quadruple mutations influence the GS/GOGAT cycle and glutathione metabolism in the cytosol. Interestingly, the content of pipecolate, which is involved in systemic acquired resistance (SAR) (Huang *et al*., 2020), was significantly increased by ∼14-fold in the quadruple mutant under −N conditions (Fig. 6b). This suggests that nitrogen deficiency strongly activates SAR in the quadruple mutant upon without pathogenic challenge.

### Decrement of basal levels of ppGpp influences phytohormone levels

We next quantified the levels of several phytohormones in WT and quadruple mutant plants grown under normal +N conditions for 12 days and then under +N and/or −N conditions for 9 days (Fig. 7). The amounts of indole-3-aceticacid (IAA), jasmonic acid (JA), and salicylic acid (SA) were higher in the quadruple mutant than in WT under +N conditions (Fig. 7a). Upon transfer to −N conditions, the WT showed no change in the levels of IAA, ABA, and SA but a reduction in the levels of JA and isoleucine-conjugated JA (JA-Ile) (Figs. 7a, b). On the other hand, the transition to −N conditions increased the levels of ABA, JA, JA-Ile, and SA in the quadruple mutant by 2–4-fold (Figs. 7a, b). Thus, the levels of ABA, JA, JA-Ile, and SA were significantly higher in the quadruple mutant than in the WT under −N conditions (Fig. 7b). These results suggest that a reduction in the basal levels of ppGpp in the quadruple mutant affects phytohormone signaling, especially under −N conditions.

**Fig. 7.**
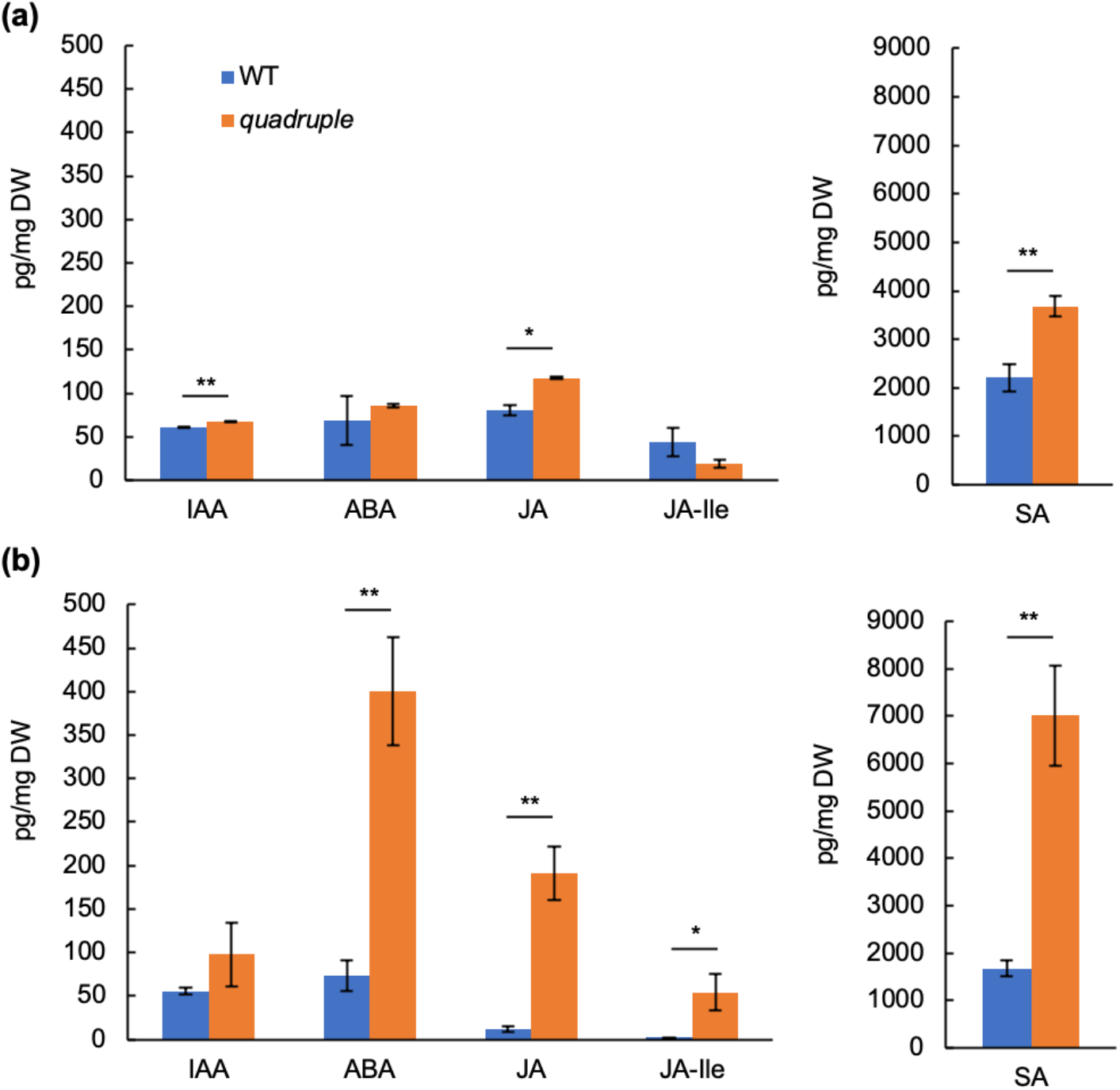
Changes in plant hormone levels in the WT and quadruple mutant under +N **(a)** and −N **(b)** conditions. After germination and growth on +N medium for 12 days, plants were transferred to +N or −N medium and sampled on day 9 for hormone analysis. Error bars represent SE (*n* = 4). Asterisks indicate significant differences (\**p* < 0.05, \*\**p* < 0.01; Student’s *t*-test). Blue: WT; Orange: quadruple mutant; ABA: abscisic acid, IAA: indole-3-acetic acid, JA: jasmonic acid, JA-Ile: jasmonic acid–isoleucine, SA: salicylic acid.

### Effects of ppGpp reduction on gene expression

To determine the influence of *rsh* quadruple mutations on the expression of plastid and nuclear genes, plants were grown under +N conditions for 14 days and then under +N and/or −N conditions for 10 days. The results of qRT-PCR revealed no significant difference among the transcript levels of seven plastid genes (*accD, clpP, rpoA, atpB, psbA, psbD*, and *rbcL*) between the WT and quadruple mutant under +N conditions (Fig. 8a). This suggests that the significant (20-fold) reduction in the basal levels of ppGpp (Fig. 2a) does not influence the transcription of plastid genes under +N conditions. Upon transition to −N conditions, the transcript levels of plastid genes increased slightly in the quadruple mutant compared with the WT, although significant difference was observed only in the expression of *rpoA*. These results suggest that ppGpp regulates plastidial gene expression under −N conditions, as reported previously (Romand *et al*., 2022).

**Fig. 8.**
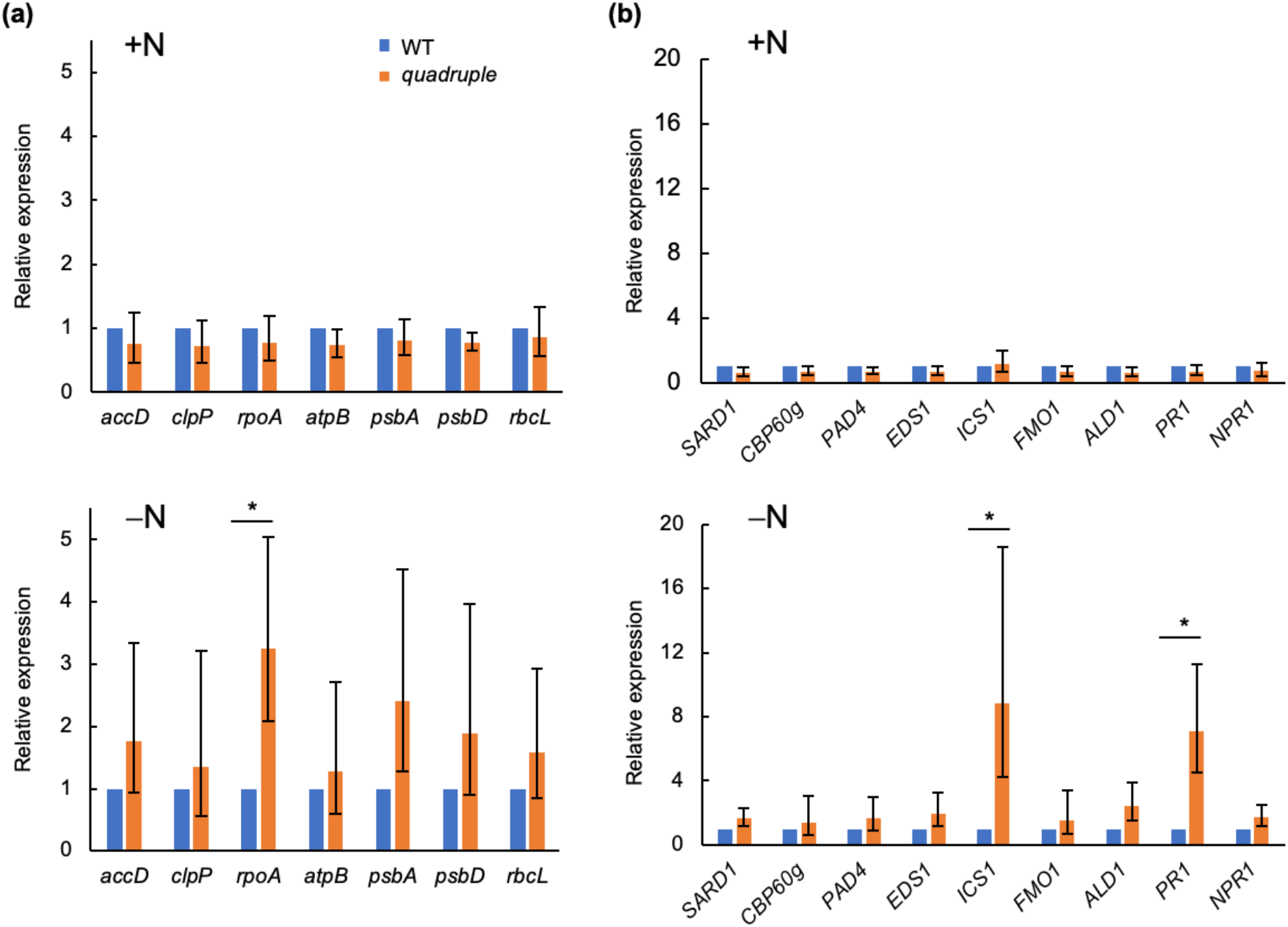
Changes in transcript levels of plastid and nuclear genes. **(a, b)** Expression levels of genes in the plastid **(a)** and nuclear **(b)** genomes of the WT and quadruple mutant under +N and −N conditions. After germination and growth on +N medium for 12 days, the plants were transferred to +N or −N medium and sampled on day 10 for RNA analysis. Expression levels in the WT were set to 1.0. Error bars represent SD (*n* = 3; \**p* < 0.05; Welch’s *t*-test).

We also analyzed the expression of nuclear defense-related genes involved in SA and/or pipecolate biosynthesis. SARD1, CBP60g, PAD4, and EDS1 are the positive regulators of isochorismate synthase ICS1, which catalyzes the conversion of chorismite to isochorismate for SA synthesis (Kim *et al*., 2020; Chen *et al*., 2020). The expression of ICS1 is a rate-limiting step for SA biosynthesis (Wildermuth *et al*., 2001; Strawn *et al*., 2007; Garcion *et al*., 2008). FMO1 and ALD1 are involved in pipecolate biosynthesis.

PR1 and NPR1 are involved in the defense response and SA perception, respectively, and their expression has been used as a marker of SA-regulated plant immune response (Wang *et al*., 2006; Kim *et al*., 2020; Chen *et al*., 2020). As shown in Fig. 8b, none of the genes tested showed any differences in expression levels between the WT and quadruple mutant under +N conditions. Under −N conditions, most genes involved in SA biosynthesis (*SARD1, CBP60g, PAD4*, and *EDS1*) and pipecolate biosynthesis (*FMO1* and *SLD1*) as well as *NPR1* showed no significant difference in expression between the quadruple mutant and WT; however, the transcript levels of *ICS1* and *PR1* were significantly higher in the quadruple mutant than in the WT.

## Discussion

To elucidate the function of the plastidial stringent response, a mutant that lacks all four RSHs has been wanted. In this study, using the CRISPR/Cas9 genome editing technology, we succeeded obtained the quadruple *rsh* mutant lacking all four *RSH* genes. The quadruple mutant showed significant (∼20-fold) reduction in basal ppGpp levels compared with the WT (Fig. 2), directly demonstrating that the four RSHs function redundantly and are responsible for maintaining the ppGpp levels in WT plastids. The source(s) of the remaining ppGpp detected in the quadruple mutant is unknown. We recently found that ppGpp is present in metazoa such as *Drosophila* as well as in human cells (Ito *et al*., 2020), suggesting that the remaining ppGpp detected in the quadruple mutant is present outside the plastids, as in metazoa, and an unknown ppGpp synthase functions in subcellular compartments other than plastids in both plants and animals. The identification of the unknown ppGpp synthetase(s) could be important for the further elucidation of ppGpp function in eukaryotes.

The ppGpp-accumulating mutant RSH3ox2, we constructed previously (Maekawa *et al*., 2015), showed higher NPQ induction than the WT under −N conditions, suggesting that the accumulated ppGpp enhances the quenching of light energy to downregulate photosynthesis (Honoki *et al*., 2018; Goto *et al*., 2022). The quadruple mutant showed lower NPQ than the WT under −N conditions (Fig. 4a), which supports the abovementioned hypothesis. Interestingly, under −N conditions, the significant reduction in Y(II) was accompanied by an increase in NPQ in RSH3ox2 (Honoki *et al*., 2018); however, the significant change in Y(II) in the quadruple mutant was not accompanied by a reduction in NPQ under −N conditions (Fig. 4). These results suggest that basal ppGpp levels under −N conditions correlate with the magnitude of NPQ induction but do not reflect photosynthetic electron transfer. This further suggests that ppGpp homeostasis, regulated redundantly by four RSHs, is required for controlled induction of NPQ in response to environmental changes. A transient increment in ppGpp level upon the transition from +N to −N conditions was recently reported (Romand *et al*., 2022), which suggests that this transient increase is also required for the photosynthetic control. To understand how ppGpp influences NPQ induction, we analyzed the expression of plastid genes by qRT-PCR. The results showed that the transcript levels of some photosynthesis-related plastid genes (Fig. 8a), as well as Y(II) values (Fig. 4b) in the quadruple mutant, were not influenced by N starvation. This suggests that the ppGpp-dependent NPQ control is not affected by the expression of photosynthetic apparatus genes, but it may be achieved at post-translation level. A recent study showed that artificial accumulation of ppGpp affects grana stacking as well as LHCII trimer aggregation in the moss *Phsycomitrella patens* (Harchouni *et al*., 2022); this probably occurred in *Arabidopsis* to induce NPQ upon the transition to −N conditions.

The GS/GOGAT cycle metabolites and glutathione metabolism were significantly altered in the quadruple mutant compared with the WT under both +N and−N conditions (Fig. 6). In WT, the level of amino acids from the GS/GOGAT cycle and shikimic acid pathway decreased and increased, respectively, upon the transition from +N to −N conditions (Honoki *et al*., 2018; Goto *et al*., 2022). Lowered metabolite levels in the GS/GOGAT cycle seem to be caused by a reduction in the Gln pool in chloroplasts (Ji *et al*., 2019). The Gln pool of chloroplasts can originate via two pathways: the transport of GS1-synthesized Gln from the cytosol to chloroplasts, and/or Gln synthesized from Glu by GS2 in chloroplasts (Perez-Garcia *et al*., 1995; Pageau *et al*., 2006; Liu *et al*., 2010). Since GS2 is sensitive to ROS (Perez-Garcia *et al*., 1995), its activity is suppressed under stress conditions, including nutrient deprivation. The *GS1* gene is upregulated under abiotic and biotic stress conditions (Liu *et al*., 2010); thus, GS1 mostly compensates for Gln depletion during the stress, suggesting that the decrement of Gln levels in the quadruple mutant is caused by the suppression of GS2 activity by the highly accumulated ROS. The ratio of GSH to GSSG relates to ROS accumulation; higher GSH:GSSG ratio results in effective ROS scavenging (Herrera-Vásquez *et al*., 2015). Metabolome analysis indicated that GSH level was 0.78-fold lower in the quadruple mutant than in the WT under +N conditions, and the GSSG level was 2-fold higher in the quadruple mutant under −N conditions (Fig. 6). This indicates that the GSH:GSSG ratio in the quadruple mutant is always lower than that in the WT, which may result in the overproduction of ROS. In fact, H_2_O_2_ levels in the quadruple mutant were higher under +N conditions, which could be eliminated in the ppGpp-overaccumulating mutant RSH3ox2 under −N conditions (Figs. 3c, d), supporting the hypothesis.

ABA, JA, JA-Ile, and SA levels were significantly higher in the quadruple mutant than in the WT, especially under −N conditions (Fig. 7). JA and JA-Ile are synthesized from α-linolenic acid, a chloroplast membrane lipid, whereas ABA is synthesized through the nonmevalonate pathway via xanthophyll carotenoids present in chloroplasts (Nambara & Marion-Poll, 2005; Mehrotra *et al*., 2014; Chini *et al*., 2018). Thus, it is reasonable to assume that the proper control of plastidial ppGpp levels is required for the biogenesis of these phytohormone. The level of H_2_O_2_ was higher in the quadruple mutant than in the WT under +N conditions (Fig. 3c), suggesting that highly production of ROS under normal conditions influences SA synthesis upon the transition to −N conditions (Noshi *et al*., 2012; Maruta *et al*., 2012; Herrera-Vásquez *et al*., 2015). The quadruple mutant showed the loss of NPQ induction upon the transition from +N to −N conditions, whereas WT plants showed an increase in NPQ (Fig. 4), which potentially contributes to ROS accumulation in the mutant. An imbalance in plastidial metabolism (Fig. 5) enhances stress responses involving JA, JA-Ile, and SA signaling, especially under −N conditions. Pipecolate accumulation in quadruple mutant plants grown under −N conditions (Fig. 6b) and imbalanced plastid metabolism caused by the accumulation of endogenous peptides and/or overproduction of ROS in chloroplasts were previously shown to induce the pathogen stress response (Kmiec *et al*., 2018; Dogra *et al*., 2022), supporting the hypothesis that imbalanced plastid metabolism is linked to the defense response.

Figure 9 illustrates a schematic model showing the role of the plastidial stringent response in the interplay between chloroplast metabolism and SA/pipecolate-dependent defense response. A certain level of ppGpp in chloroplasts is required for proper metabolism. Upon N starvation, the ppGpp-dependent metabolic control, as well as NPQ induction in chloroplasts, suppressed excess ROS accumulation, which was impaired in the quadruple mutant. Additionally, ppGpp maintained the homeostasis of the GSH:GSSG ratio in the WT to suppress ROS accumulation; this was also impaired in *quadruple* mutant. Consequently, SA and pipecolate levels increased in the quadruple mutant, but not in the WT, under −N conditions.

**Fig. 9.**
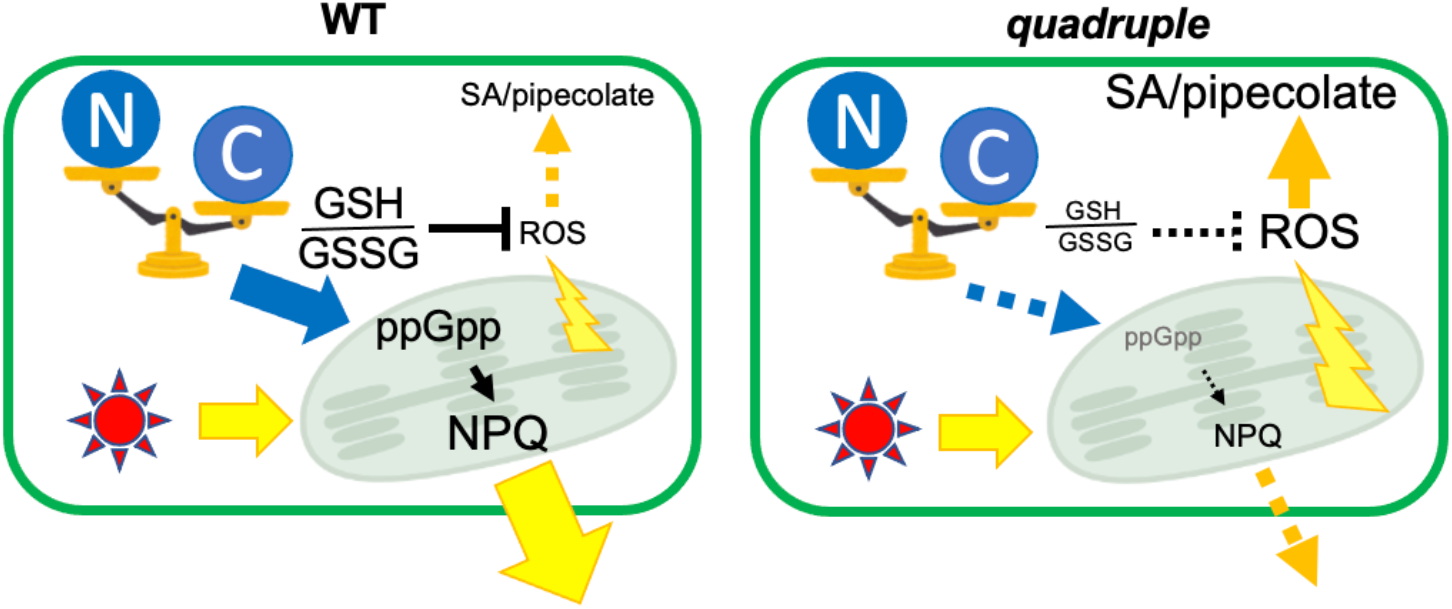
Schematic model of the role of the plastidial stringent response in ROS accumulation and plant defense response.

In summary, significant decrease in the basal levels of ppGpp by quadruple *rsh* mutations influenced plastidial metabolism and the loss of NPQ induction upon −N transition, indicating the crucial role of the plastidial stringent response for C/N balance response. Under −N conditions, the quadruple mutant accumulated SA and pipecolate, which are involved in the plant defense response. This indicates that the control of plastidial metabolism by ppGpp is required for the interplay between nitrogen starvation and pathogen stress response. The crucial role of chloroplast metabolism in the plant defense response has long been discussed, although detailed mechanisms remain elusive (Kangasjärvi *et al*., 2012; Trotta *et al*., 2014; Delprato *et al*., 2015; Roeber *et al*., 2021). Further research using the *rsh*-quadruple mutant, constructed in this study, is needed to uncover the functional role of the plastidial stringent response in phytohormone signaling and plant defense response under nutrient limiting conditions.

## Materials and Methods

### Plant materials and growth conditions

*Arabidopsis thaliana* (ecotype Columbia) and its mutants were grown on half-strength Murashige and Skoog (1/2 MS) medium, containing 0.8% agar and 1% (w/v) sucrose, at 23°C under continuous light conditions (40 µmol photons m^−2^ s^−1^). To prepare medium for the nitrogen starvation treatment, KNO_3_ in 1/2 MS medium was replaced by KCl at the same concentration, and NH_4_NO_3_ was removed, and 0.1 mM KNO_3_ and 0.1 mM NH_4_NO_3_ were added. To generate the *rsh1 rsh2 rsh3 crsh* quadruple mutant, the *CRSH* gene in the *rsh1 rsh2 rsh3* triple mutant (Ono *et al*., 2021) was mutated using the CRISPR/Cas9 technology (Fauser *et al*., 2014). Briefly, a protospacer sequence was designed in the first exon of *CRSH* (83–102 bp from the translation start site), and the protospacer oligonucleotides were annealed and cloned into the pEn-Chimera plasmid, as described previously (Fauser *et al*., 2014). After confirming its sequence, the DNA insert was further cloned into pDe-CAS9 using Gateway LR Clonase (ThermoFisher, Waltham, Massachusetts, USA). The obtained plasmid was introduced into the wild type (WT) with the standard floral dip method using *Agrobacterium* to induce double-strand breaks at the protospacer position. Homozygous mutations in the quadruple mutant were confirmed by PCR using gene-specific primers (Ono *et al*., 2021), followed by sequencing. RNA was isolated using the SV Total RNA Isolation System (Promega, Madison, Wisconsin, USA). Then, first-strand cDNA was synthesized with the PrimeScript RT Reagent Kit (TaKaRa, Shiga, Japan), and genomic DNA isolated from the WT. Subsequently, RT-PCR was performed with gene-specific primers using the cDNA and genomic DNA as templates.

### Quantitative mRNA analysis

Total RNA was isolated from WT and mutant plants grown on 1/2 MS medium under continuous light (40 µmol photons m^−2^ s^−1^) for 14 d, as described previously (Goto *et al*., 2022). The 14-day-old plants were transferred to normal and/or nitrogen-starvation medium, as described above, and grown on each medium for 10 days. Then, the plants were harvested and subjected to total RNA isolation using the SV Total RNA Isolation System (Promega, Madison, Wisconsin, USA). First-strand cDNA was synthesized with the PrimeScript RT Reagent Kit (TaKaRa, Shiga, Japan), and qRT-PCR was performed on the Thermal Cycler Dice® (TaKaRa) using SYBR® Premix EX Taq™ and gene-specific primers designed in this study (Table 2) or previously (for plastidial genes; Sano *et al*., 2014; Maekawa *et al*., 2015; Kim *et al*., 2020). The *Ubiquitin 10* (*UBQ10*) gene, which shows a constant expression level independent of the growth conditions (Sun & Callis, 1997), was used as an internal standard.

**Table 2.**
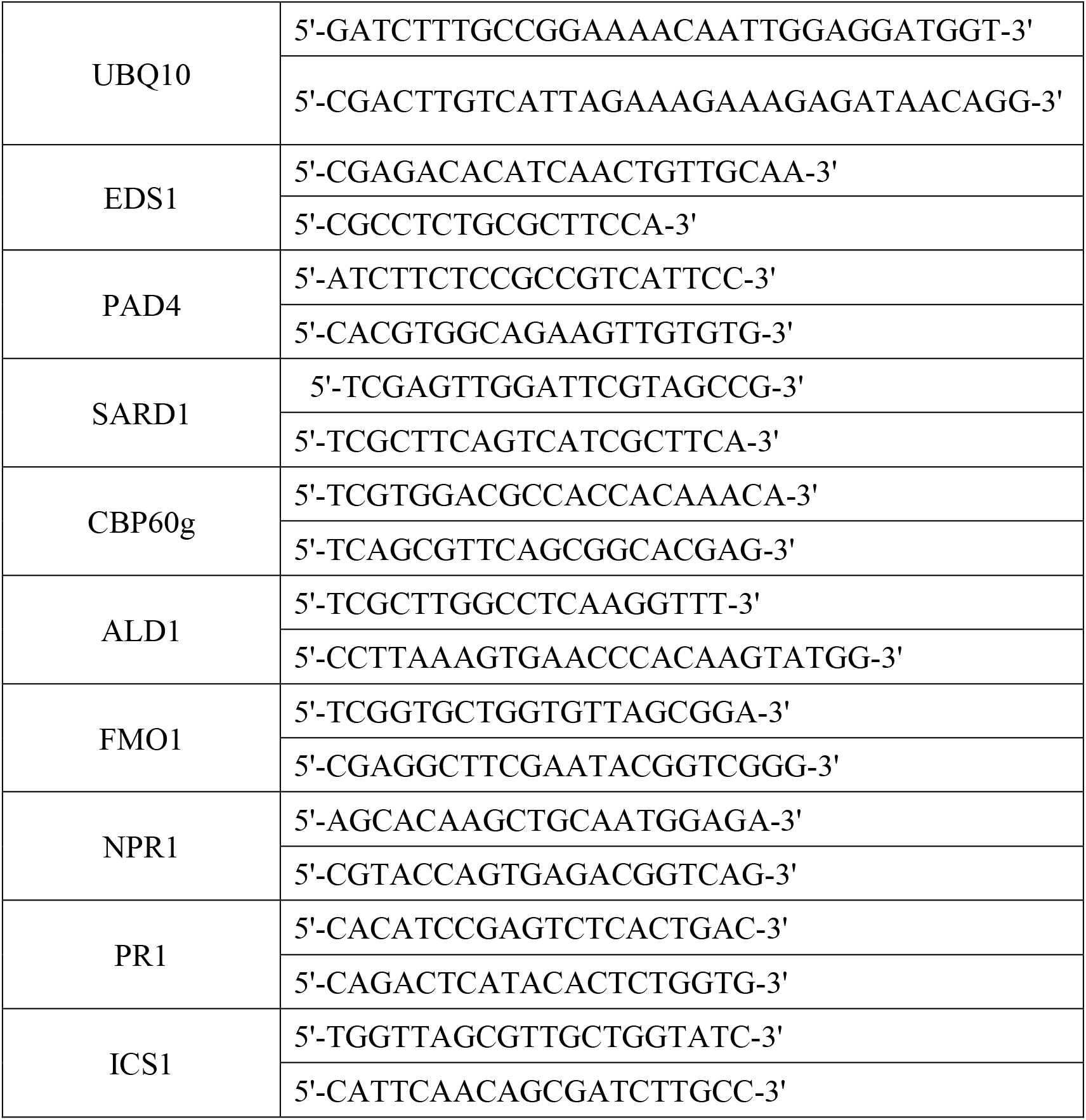
Primers used in this study

### Measurement of chlorophyll, carotenoid, ppGpp, metabolite, and phytohormone contents

Plants were grown on solid 1/2 MS medium, containing 0.8% agar, under continuous light (40 µmol photons m^−2^ s^−1^). Pigments were extracted and quantified as described previously (Goto *et al*., 2022). Chlorophyll (Chl) *a* and *b* and carotenoids (Car) were quantified as described by Lichtenthaler (1987) and Porra *et al*. (1989). The extraction and quantification of ppGpp were performed as described by Ihara *et al*. (2015). The metabolome analysis of plants was performed as described by Oikawa *et al*. (2011). Raw metabolome data were subjected to Partial Least-Squares Discriminant Analysis (PLS-DA) on MetaboAnalyst 5.0 (https://www.metaboanalyst.ca. Seven metabolites including stachydrine, Ala–Ala, phenylpyruvate, galacturonate; glucuronate, deoxyinosine, glycoshenodeoxycholate, and NeuGc, were omitted as they showed identical and/or no values in some individual datasets. Analysis was performed with default settings in the autoscaling mode. Plant hormone analysis was performed as described by Kanno *et al*. (2016).

### Hydrogen peroxide (H_2_O_2_) quantification

The amount of H_2_O_2_ in shoots was measured as described previously (Junglee *et al*., 2014), with slight modifications. Briefly, frozen shoot samples were homogenized with 2.5 mM potassium phosphate buffer (pH 7.0), 0.05% trichloroacetic acid (TCA; w/v), and 250 mM potassium iodide, and then incubated in the dark for 10 min. The samples were transferred to new tubes, and centrifuged at 12,000 × *g* for 15 min at 4 °C. Then, the supernatant of each sample was transferred to a new tube in the dark, and absorbance was measured at 390 nm using the SHIMAZU UV-1800 spectrophotometer. A standard curve was generated using serial dilutions of H_2_O_2_ (FUJIFILM-WAKO Chemicals, Osaka, Japan). The quantity of H_2_O_2_ in each sample was normalized relative to the sample fresh weight.

### Chl fluorescence measurements

Plants were incubated in the dark for 30 min. Then, Chl fluorescence parameters were measured under actinic light (55 μmol photons m^−2^ s^−1^) using the Dual-PAM-100 system (Walz, Effeltrich, Germany). The maximum quantum yield pf photosystem II (PSII; Fv/Fm) was calculated as (Fm−Fo)/Fm. The effective quantum yield of PSII (Y[II]) during steady-state photosynthesis was calculated as (Fm′−Fs)/Fm′. Nonphotochemical quenching (NPQ) was calculated as (Fm−Fm′)/Fm′.

### Western blot analysis

Protein amounts were determined using the Bradford Assay Kit (Bio-Rad, Hercules, California, USA), and 6.0 μg of each protein was separated by sodium dodecyl sulfate-polyacrylamide gel electrophoresis (SDS-PAGE). The SDS-PAGE gels were electroblotted onto polyvinylidene difluoride (PVDF) membranes (GE-Healthcare, Chicago, Illinois, USA), which were then incubated with anti-CRSH antibody (Masuda *et al*., 2008). Immunoreactive proteins were detected using the Alkaline Phosphatase Substrate Kit II (Vector Laboratories, Burlingame, California, USA). Membranes were stained with Coomassie brilliant blue to visualize the proteins.

### Statistical analysis

All experiments were designed and performed with at least three independent replicates. Calculations were performed and graphs and charts were constructed using Microsoft Excel and the *R* software. Statistical differences among the experimental samples were confirmed using Student’s *t*-test, Welch’s *t*-test, Dunnett test, or Turkey test with 95% confidence intervals. Other details such as sample sizes are shown in each legend.

## Abbreviations

CRSH: Ca^2+^-activated
RSH: ppGpp, 5’-diphosphate 3’-diphosphate;
RSH: RelA-SpoT homolog

## Acknowledgements

We thank the Biomaterial Analysis Division of Tokyo Institute of Technology for support with devices used and technical assistance. This work was supported by a Grant-in-Aid for Scientific Research, KAKENHI (21H02075) to SM.

## References

Abdelkefi, H, Sugliani M, Ke H, Harchouni S, Soubigou-Taconnat L, Citerne S, Mouille G, Fakhfakh H, Robaglia C, Field B. 2018. Guanosine tetraphosphate modulates salicylic acid signalling and the resistance of Arabidopsis thaliana to Turnip mosaic virus. Molecular Plant Pathology 19: 634–646.

Avilan L, Lebrun R, Puppo C, Citerne S, Cuiné S, Li-Beisson Y, Menand B, Field B, Gontero B. 2021. ppGpp influences protein protection, growth and photosynthesis in Phaeodactylum tricornutum. The New phytologist 230: 1517–1532.

Avilan L, Puppo C, Villain A, Bouveret E, Menand B, Field B, Gontero B. 2019. RSH enzyme diversity for (p)ppGpp metabolism in Phaeodactylum tricornutum and other diatoms. Scientific reports 9.

van der Biezen EA, Sun J, Coleman MJ, Bibb MJ, Jones JDG. 2000. Arabidopsis RelA/SpoT homologs implicate (p)ppGpp in plant signaling. Proceedings of the National Academy of Sciences 97: 3747–3752.

Cashel M. 1969. The control of ribonucleic acid synthesis in Escherichia coli IV. Relevance of unusual phosphorylated compounds from amino acid-starved stringent strains. J. Biol. Chem. 244: 3133–3141.

Cashel M, Gentry DR, Hernandez VJ, Vinella D. 1996. The stringent response. In: Neidhardt FC, Curtiss IR, Ingraham JL, Lin Ecc, Low KB, Magasanik B, Reznikoff WS, Riley M, Schaechter M, Umbarger HE, eds. Escherichia coli and Salmonella: cellular and molecular biology. Washington D.C.: ASM Press, 1458–1496.

Chen J, Clinton M, Qi G, Wang D, Liu F, Qing Fu Z. 2020. Reprogramming and remodeling: transcriptional and epigenetic regulation of salicylic acid-mediated plant defense. Journal of Experimental Botany 71: 5256–5268.

Chini A, Monte I, Zamarreño AM, Hamberg M, Lassueur S, Reymond P, Weiss S, Stintzi A, Schaller A, Porzel A, et al. 2018. An OPR3-independent pathway uses 4,5-didehydrojasmonate for jasmonate synthesis. Nature Chemical Biology 2018 14:2 14: 171–178.

Delprato ML, Krapp AR, Carrillo N. 2015. Green Light to Plant Responses to Pathogens: The Role of Chloroplast Light-Dependent Signaling in Biotic Stress. Photochemistry and Photobiology 91: 1004–1011.

Dogra V, Singh RM, Li M, Li M, Singh S, Kim C. 2022. EXECUTER2 modulates the EXECUTER1 signalosome through its singlet oxygen-dependent oxidation. Molecular Plant 15: 438–453.

Fauser F, Schiml S, Puchta H. 2014. Both CRISPR/Cas-based nucleases and nickases can be used efficiently for genome engineering in Arabidopsis thaliana. The Plant Journal 79: 348–359.

Field B. 2018. Green magic: regulation of the chloroplast stress response by (p)ppGpp in plants and algae. Journal of Experimental Botany 69: 2797–2807.

Garcion C, Lohmann A, Lamodière E, Catinot J, Buchala A, Doermann P, Métraux JP. 2008. Characterization and biological function of the ISOCHORISMATE SYNTHASE2 gene of Arabidopsis. Plant physiology 147: 1279–1287.

Givens RM, Lin M-H, Taylor DJ, Mechold U, Berry JO, Hernandez VJ. 2004. Inducible expression, enzymatic activity, and origin of higher plant homologues of bacterial RelA/SpoT stress proteins in Nicotiana tabacum. The Journal of biological chemistry 279: 7495–504.

Goto M, Oikawa A, Masuda S. 2022. Metabolic changes contributing to large biomass production in the Arabidopsis ppGpp-accumulating mutant under nitrogen deficiency. Planta 255.

Harchouni S, England S, Vieu J, Aouane A, Citerne S, Legeret B, Li-Beisson Y, Menand B, Field B. 2022. Guanosine tetraphosphate (ppGpp) accumulation inhibits chloroplast gene expression and promotes super grana formation in the moss Physcomitrium (Physcomitrella) patens. New Phytol.

Herrera-Vásquez A, Salinas P, Holuigue L. 2015. Salicylic acid and reactive oxygen species interplay in the transcriptional control of defense genes expression. Frontiers in Plant Science 6: 1–9.

Honoki R, Ono S, Oikawa A, Saito K, Masuda S. 2018. Significance of accumulation of the alarmone (p)ppGpp in chloroplasts for controlling photosynthesis and metabolite balance during nitrogen starvation in Arabidopsis. Photosynthesis Research 135: 299– 308.

Huang W, Wang Y, Li X, Zhang Y. 2020. Biosynthesis and Regulation of Salicylic Acid and N-Hydroxypipecolic Acid in Plant Immunity. Molecular Plant 13: 31–41.

Ihara Y, Ohta H, Masuda S. 2015. A highly sensitive quantification method for the accumulation of alarmone ppGpp in Arabidopsis thaliana using UPLC-ESI-qMS/MS. Journal of Plant Research 128.

Imamura S, Nomura Y, Takemura T, Pancha I, Taki K, Toguchi K, Tozawa Y, Tanaka K. 2018. The checkpoint kinase TOR (target of rapamycin) regulates expression of a nuclear-encoded chloroplast RelA-SpoT homolog (RSH) and modulates chloroplast ribosomal RNA synthesis in a unicellular red alga. The Plant Journal 94: 327–339.

Ito D, Ihara Y, Nishihara H, Masuda S. 2017. Phylogenetic analysis of proteins involved in the stringent response in plant cells. J. Plant Res. 130: 625–634.

Ito K, Ito D, Goto M, Suzuki S, Masuda S, Iba K, Kusumi K. 2022. Regulation of ppGpp Synthesis and Its Impact on Chloroplast Biogenesis during Early Leaf Development in Rice. Plant and Cell Physiology.

Ito D, Kawamura H, Oikawa A, Ihara Y, Shibata T, Nakamura N, Asano T, Kawabata S-I, Suzuki T, Masuda S. 2020. ppGpp functions as an alarmone in metazoa. Communications Biology 2020 3:1 3: 1–11.

Ji Y, Li Q, Liu G, Selvaraj G, Zheng Z, Zou J, Wei Y. 2019. Roles of Cytosolic Glutamine Synthetases in Arabidopsis Development and Stress Responses. Plant and Cell Physiology 60: 657–671.

Junglee S, Urban L, Sallanon H, Lopez-Lauri F, Junglee S, Urban L, Sallanon H, Lopez-Lauri F. 2014. Optimized Assay for Hydrogen Peroxide Determination in Plant Tissue Using Potassium Iodide. American Journal of Analytical Chemistry 5: 730–736.

Kangasjärvi S, Neukermans J, Li S, Aro EM, Noctor G. 2012. Photosynthesis, photorespiration, and light signalling in defence responses. Journal of Experimental Botany 63: 1619–1636.

Kanno Y, Oikawa T, Chiba Y, Ishimaru Y, Shimizu T, Sano N, Koshiba T, Kamiya Y, Ueda M, Seo M. 2016. AtSWEET13 and AtSWEET14 regulate gibberellin-mediated physiological processes. Nature Communications 7: 13245.

Kim Y, Gilmour SJ, Chao L, Park S, Thomashow MF. 2020. Arabidopsis CAMTA Transcription Factors Regulate Pipecolic Acid Biosynthesis and Priming of Immunity Genes. Molecular Plant 13: 157–168.

Kmiec B, Branca RMM, Berkowitz O, Li L, Wang Y, Murcha MW, Whelan J, Lehtiö J, Glaser E, Teixeira PF. 2018. Accumulation of endogenous peptides triggers a pathogen stress response in Arabidopsis thaliana. The Plant Journal 96: 705–715.

Li H, Nian J, Fang S, Guo M, Huang X, Zhang F, Wang Q, Zhang J, Bai J, Dong G, et al. 2022. Regulation of nitrogen starvation responses by the alarmone (p)ppGpp in rice. Journal of Genetics and Genomics 49: 469–480.

Lichtenthaler HK. 1987. Chlorophylls and carotenoids: Pigments of photosynthetic biomembranes. Methods in Enzymology 148: 350–382.

Liu G, Ji Y, Bhuiyan NH, Pilot G, Selvaraj G, Zou J, Wei Y. 2010. Amino Acid Homeostasis Modulates Salicylic Acid–Associated Redox Status and Defense Responses in Arabidopsis. The Plant Cell 22: 3845–3863.

Maekawa M, Honoki R, Ihara Y, Sato R, Oikawa A, Kanno Y, Ohta H, Seo M, Saito K, Masuda S. 2015. Impact of the plastidial stringent response in plant growth and stress responses. Nature Plants 1: 15167.

Maruta T, Noshi M, Tanouchi A, Tamoi M, Yabuta Y, Yoshimura K, Ishikawa T, Shigeoka S. 2012. H2O2-triggered Retrograde Signaling from Chloroplasts to Nucleus Plays Specific Role in Response to Stress. Journal of Biological Chemistry 287: 11717– 11729.

Masuda S. 2012. The stringent response in phototrophs. In: Najafpour M, ed. Advances in photosynthesis. In Tech., 487–500.

Masuda S, Mizusawa K, Narisawa T, Tozawa Y, Ohta H, Takamiya K-I. 2008. The bacterial stringent response, conserved in chloroplasts, controls plant fertilization. Plant & cell physiology 49: 135–41.

Mehrotra R, Bhalothia P, Bansal P, Basantani MK, Bharti V, Mehrotra S. 2014. Abscisic acid and abiotic stress tolerance – Different tiers of regulation. Journal of Plant Physiology 171: 486–496.

Mizusawa K, Masuda S, Ohta H. 2008. Expression profiling of four RelA/SpoT-like proteins, homologues of bacterial stringent factors, in Arabidopsis thaliana. Planta 228: 553–62.

Nambara E, Marion-Poll A. 2005. ABSCISIC ACID BIOSYNTHESIS AND CATABOLISM. Annu. Rev. Plant Biol. 56: 165–185.

Noshi M, Maruta T, Shigeoka S. 2012. Relationship between chloroplastic H2O2 and the salicylic acid response. https://doi.org/10.4161/psb.20906 7: p944–946.

Oikawa A, Fujita N, Horie R, Saito K, Tawaraya K. 2011. Solid-phase extraction for metabolomic analysis of high-salinity samples by capillary electrophoresis-mass spectrometry. J. Sep. Sci. 34: 1063–1068.

Ono S, Suzuki S, Ito D, Tagawa S, Shiina T, Masuda S. 2021. Plastidial (p)ppGpp Synthesis by the Ca2+-Dependent RelA-SpoT Homolog Regulates the Adaptation of Chloroplast Gene Expression to Darkness in Arabidopsis. Plant & Cell Physiology 61: 2077–2086.

Pageau K, Reisdorf-Cren M, Morot-Gaudry JF, Masclaux-Daubresse C. 2006. The two senescence-related markers, GS1 (cytosolic glutamine synthetase) and GDH (glutamate dehydrogenase), involved in nitrogen mobilization, are differentially regulated during pathogen attack and by stress hormones and reactive oxygen species in Nicotiana tabacum L. leaves. Journal of Experimental Botany 57: 547–557.

Perez-Garcia A, Canovas FM, Gallardo F, Hirel B, de Vicente A. 1995. Differential expression of glutamine synthetase isoforms in tomato detached leaflets infected with Pseudomonas syringae pv. tomato. Molecular Plant-Microbe Interactions 8: 96–103.

Porra R., Thompson W., Kriedemann P. 1989. Determination of accurate extinction coefficients and simultaneous equations for assaying chlorophylls a and b extracted with four different solvents: verification of the concentration of chlorophyll standards by atomic absorption spectroscopy. Biochim. Biophys. Acta 975: 384–394.

Potrykus K, Cashel M. 2008. (p)ppGpp: Still Magical? *. Annual Review of Microbiology 62: 35–51.

Roeber VM, Bajaj I, Rohde M, Schmülling T, Cortleven A. 2021. Light acts as a stressor and influences abiotic and biotic stress responses in plants. Plant, Cell & Environment 44: 645–664.

Romand S, Abdelkefi H, Lecampion C, Belaroussi M, Dussenne M, Ksas B, Citerne S, Caius J, D’alessandro S, Fakhfakh H, et al. 2022. A guanosine tetraphosphate (ppGpp) mediated brake on photosynthesis is required for acclimation to nitrogen limitation in Arabidopsis. eLife 11.

Sano S, Aoyama M, Nakai K, Shimotani K, Yamasaki K, Sato MH, Tojo D, Suwastika IN, Nomura H, Shiina T. 2014. Light-dependent expression of flg22-induced defense genes in Arabidopsis. Frontiers in Plant Science 5: 531.

Strawn MA, Marr SK, Inoue K, Inada N, Zubieta C, Wildermuth MC. 2007. Arabidopsis isochorismate synthase functional in pathogen-induced salicylate biosynthesis exhibits properties consistent with a role in diverse stress responses. The Journal of biological chemistry 282: 5919–5933.

Sugliani M, Abdelkefi H, Ke H, Bouveret E, Robaglia C, Caffarri S, Field B. 2016. An ancient bacterial signaling pathway regulates chloroplast function to influence growth and development in Arabidopsis. Plant Cell 28: 661–679.

Sun CW, Callis J. 1997. Independent modulation of Arabidopsis thaliana polyubiquitin mRNAs in different organs and in response to environmental changes. The Plant journal : for cell and molecular biology 11: 1017–1027.

Takahashi K, Kasai K, Ochi K. 2004. Identification of the bacterial alarmone guanosine 5’-diphosphate 3’-diphosphate (ppGpp) in plants. Proc. Natl. Acad. Sci. USA 101: 4320–4324.

Tozawa Y, Nomura Y. 2011. Signalling by the global regulatory molecule ppGpp in bacteria and chloroplasts of land plants. Plant biology (Stuttgart, Germany) 13: 699– 709.

Tozawa Y, Nozawa A, Kanno T, Narisawa T, Masuda S, Kasai K, Nanamiya H. 2007. Calcium-activated (p)ppGpp synthetase in chloroplasts of land plants. The Journal of biological chemistry 282: 35536–45.

Trotta A, Rahikainen M, Konert G, Finazzi G, Kangasjärvi S. 2014. Signalling crosstalk in light stress and immune reactions in plants. Philosophical Transactions of the Royal Society B: Biological Sciences 369.

Wang D, Amornsiripanitch N, Dong X. 2006. A genomic approach to identify regulatory nodes in the transcriptional network of systemic acquired resistance in plants. PLoS pathogens 2: 1042–1050.

Wildermuth MC, Dewdney J, Wu G, Ausubel FM. 2001. Isochorismate synthase is required to synthesize salicylic acid for plant defence. Nature 414: 562–565.

Yamburenko M V., Zubo YO, Börner T. 2015. Abscisic acid affects transcription of chloroplast genes via protein phosphatase 2C-dependent activation of nuclear genes: repression by guanosine-3′-5′-bisdiphosphate and activation by sigma factor 5. The Plant Journal 82: 1030–1041.

